# Evolution of Opsin Genes in Caddisflies (Insecta: Trichoptera)

**DOI:** 10.1101/2023.12.15.571940

**Authors:** Ashlyn Powell, Jacqueline Heckenhauer, Steffen U. Pauls, Blanca Ríos-Touma, Ryoichi B. Kuranishi, Ralph W. Holzenthal, Ernesto Razuri-Gonzales, Seth Bybee, Paul B. Frandsen

## Abstract

Insects have evolved complex and diverse visual systems in which light-sensing protein molecules called opsins couple with a chromophore to form photopigments. Insect photopigments group into three major gene families based on wavelength sensitivity: long wavelength (LW), short wavelength (SW), and ultraviolet wavelength (UV). Here, we identified 123 opsin sequences from whole genome assemblies across 25 caddisfly species (Insecta: Trichoptera). We discovered the LW opsins have the most diversity across species and form two separate clades in the opsin gene tree. Conversely, we observed a loss of the SW opsin in half of the trichopteran species in this study, which might be associated with the fact that caddisflies are active during low-light conditions. Lastly, we found a single copy of the UV opsin in all the species in this study, with one exception: *Athripsodes cinereus* has two copies of the UV opsin and resides within a clade of caddisflies with colorful wing patterns.

**Significance:** While opsin evolution in some insect groups has been well-characterized, it has never been studied across caddisflies. Our findings provide insight into the diversity of opsins in caddisflies and form a basis for further research into the evolutionary drivers and complex visual systems in Trichoptera.

## Introduction

Within the visual system, the ability to perceive light is critical and plays an essential role in the life histories of insects including finding food, avoiding predators, and selecting a mate (van der Kooi et al., 2021). Light perception occurs primarily within three different types of visual organs in insects: the stemmata of larvae and the ocelli and compound eyes of adults (Guignard et al., 2022; van der Kooi et al., 2021). Upon light absorption, photoreceptors within the eyes—which contain opsin proteins and chromophores—change their configuration from a resting state to a signaling state, thereby indicating a physiological response (Shichida & Matsuyama, 2009). Insect visual opsins form three major gene clades based on their peak wavelength sensitivity, including long wavelength (LW, 500–600 nm), short wavelength (SW, 400–500 nm), and ultraviolet wavelength (UV, 300–400 nm) (Feuda et al., 2016; Lord et al., 2016, van der Kooi et al., 2021). Many insect groups possess an additional opsin type, Rhodopsin 7 (RH7), which does not have a known function in most insect groups but was found to be involved in circadian rhythms in *Drosophila* (Ni et al., 2017; Senthilan & Helfrich-Förster, 2016).

Insects typically possess one or more copies of each opsin type. Moreover, multiple cases of gene duplications and losses have been observed throughout insect opsin evolution (Briscoe et al., 2010; Feuda et al., 2016; French et al., 2015; Frentiu et al., 2007; Friedrich, 2023; Futahashi et al., 2015; Guignard et al., 2022; Lord et al., 2016; Mulhair et al., 2023; Sharkey et al., 2017; Sison-Mangus et al., 2008; Sondhi et al., 2021; Spaethe and Briscoe, 2004; Suvorov et al., 2017). These duplications, and subsequent diversification, of visual opsin genes are the primary mechanisms of evolution that lead to greater visual capacity and flexibility (Frentiu et al., 2007; Friedrich, 2023; Suvorov et al., 2017) and are usually linked to a particular life history strategy, living environment, or light condition (French et al., 2015; Futahashi et al., 2015; Guignard et al., 2022; Lord et al., 2016; Sharkey et al., 2017; 2023; Sondhi et al., 2021). While the evolution of opsins has been relatively well studied in some insect orders, opsin genes in caddisflies (Insecta: Trichoptera) have only been characterized in a single species as part of a broader comparative study across insects (Guignard et al., 2022).

As eggs, larvae, and pupae, caddisflies mainly inhabit the benthic zone of freshwater habitats, but as adults, they occupy terrestrial environments adjacent to freshwater (Morse et al., 2019). There are two monophyletic suborders within Trichoptera characterized by differences in habitat, morphology, and silk use: Annulipalpia (retreat-making) and Integripalpia (cocoon– and case-making) (Frandsen et al., 2024). Adult caddisflies resemble small moths; yet, while most species have wings and bodies covered in small hairs instead of scales, a few species have brightly colored wings with red, orange, green, or silver regions (Fig. 1) due to the development of hairs into scales (Holzenthal et al., 2007). Additionally, adult caddisflies possess varying eye sizes, some much larger than others (Fig. 1). Presumably, such varied environments—both aquatic and terrestrial—and wing colorations require a plastic and diverse visual system. To gain an understanding of opsin evolution in caddisflies, we analyzed the occurrence and phylogenetic relationships of opsin genes from whole genome assemblies across 25 caddisfly species (Table 1), representing the major evolutionary lineages within the order.

**Figure 1:**
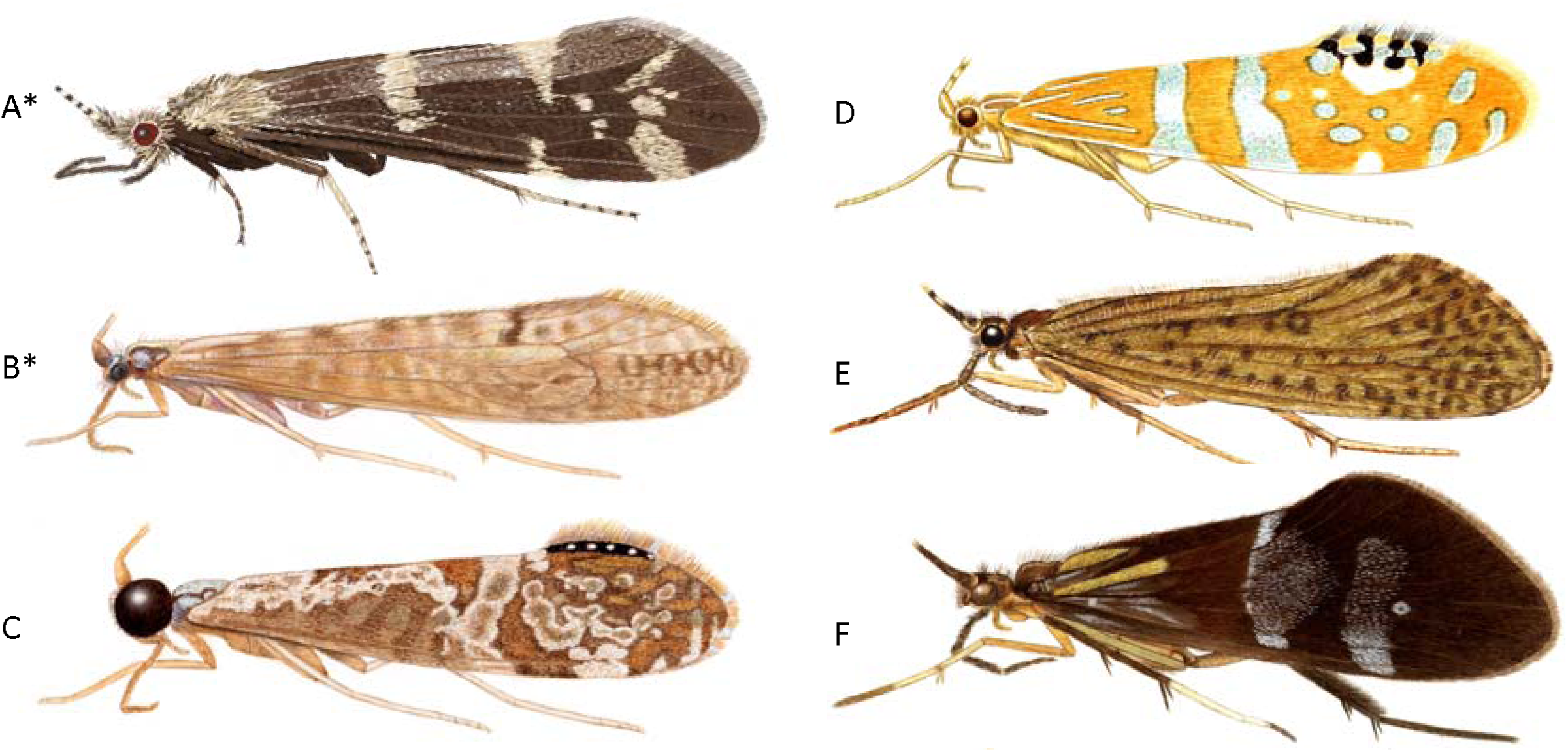
Adult caddisfly illustrations showing varying eye sizes and diverse wing colorations and patterns. (A) *Athripsodes cinereus* (Leptoceridae); (B) *Nectopsyche paramo* (Leptoceridae); (C) *Nectopsyche nigricapilla* (Leptoceridae); (D) *Nectopsyche ortizi* (Leptoceridae); (E) *Banyallarga vicaria* (Calamoceratidae); (F) *Phylloicus abdominalis* (Calamoceratidae). (*) Species included in this study. Illustrations by Julie Martinez and Ralph Holzenthal.

**Table 1:** Locations and quality scores of genome assemblies for each species. Compleasm (BUSCO) was run with the Endopterygota OrthoDB v10 gene set and the results categories are as follows: C: complete; S: single; D: duplicated; F: fragmented subclass 1 (only a portion of the gene is present in the assembly); I: fragmented subclass 2 (different sections of the gene align to different locations in the assembly); M: missing.

| Species | Accession number | Publication | Contig N50 (bp) | Compleasm (BUSCO) |
| --- | --- | --- | --- | --- |
| *Leptonema lineaticorne* | GCA_024500535.1 | Heckenhauer et al. 2023 | 14,931,587 | C:98.30%[S:97.83%,D:0.47%],<br>F:0.66%,I:0.00%,M:1.04% |
| *Hydropsyche tenuis* | GCA_009617725.1 | Heckenhauer et al. 2019 | 2,190,134 | C:97.93%[S:97.65%,D:0.28%],<br>F:0.89%,I:0.00%,M:1.18% |
| *Parapsyche elsis* | GCA_022651745.1 | Frandsen et al. 2019,<br>Heckenhauer et al. 2022 | 5,591,679 | C:96.99%[S:96.80%,D:0.19%],<br>F:0.80%,I:0.00%,M:2.21% |
| *Arctopsyche grandis* | GCA_029955255.1 | Frandsen et al. 2023 | 6,470,670 | C:98.36%[S:95.39%,D:2.97%],<br>F:0.71%,I:0.00%,M:0.94% |
| *Plectrocnemia conspersa* | Available on FigShare | New | 32,103,979 | C:97.74%[S:97.32%,D:0.42%],<br>F:0.61%,I:0.00%,M:1.65% |
| *Philopotamus ludificatus* | GCA_022495035.1 | Heckenhauer et al. 2022 | 35,449 | C:94.63%[S:93.31%,D:1.32%],<br>F:3.15%,I:0.05%,M:2.17% |
| *Stenopsyche tienmushanensis* | GCA_008973525.1 | Luo et al. 2018 | 1,296,863 | C:97.50%[S:95.15%,D:2.35%],<br>F:1.04%,I:0.00%,M:1.46% |
| *Agraylea sexmaculata* | GCA_022606485.1 | Heckenhauer et al. 2022 | 86,524 | C:95.67%[S:90.96%,D:4.71%],<br>F:1.04%,I:0.00%,M:3.30% |
| *Glossosoma conforme* | GCA_022606575.1 | Heckenhauer et al. 2022 | 2,212,131 | C:93.13%[S:92.47%,D:0.66%],<br>F:0.66%,I:0.00%,M:6.21% |
| *Atopsyche davidsoni* | GCA_022113835.1 | Ríos-Touma et al. 2021 | 14,095,054 | C:98.68%[S:98.35%,D:0.33%],<br>F:0.61%,I:0.00%,M:0.71% |
| *Atopsyche callosa* | Available on FigShare | New | 25,586,909 | C:98.54%[S:96.70%,D:1.84%],<br>F:0.66%,I:0.00%,M:0.80% |
| *Himalopsyche phryganea* | GCA_022494535.1 | Heckenhauer et al. 2019 | 4,634,010 | C:98.21%[S:97.83%,D:0.38%],<br>F:0.71%,I:0.00%,M:1.08% |
| *Himalopsyche tibetana* | GCA_030503985.1 | Heckenhauer et al. 2022 | 28,889,006 | C:98.91%[S:98.35%,D:0.56%],<br>F:0.56%,I:0.00%,M:0.52% |
| *Rhyacophila brunnea* | Available on FigShare | New | 1,306,779 | C:97.74%[S:94.35%,D:3.39%],<br>F:1.22%,I:0.00%,M:1.04% |
| *Micrasema minimum* | GCA_022494985.1 | Heckenhauer et al. 2019 | 69,526 | C:66.34%[S:66.20%,D:0.14%],<br>F:7.25%,I:0.00%,M:26.41% |
| *Lepidostoma basale* | GCA_022606425.2 | Heckenhauer et al. 2019 | 1,001,566 | C:98.16%[S:97.55%,D:0.61%],<br>F:0.85%,I:0.05%,M:0.94% |
| *Drusus annulatus* | GCA_022651775.1 | Heckenhauer et al. 2019 | 1,032,046 | C:97.92%[S:97.50%,D:0.42%],<br>F:1.08%,I:0.00%,M:0.99% |
| *Halesus radiatus* | GCA_022606495.2 | Heckenhauer et al. 2019 | 124,280 | C:91.20%[S:89.69%,D:1.51%],<br>F:4.38%,I:0.05%,M:4.38% |
| *Hesperophylax magnus* | GCA_026573805.1 | Frandsen et al. 2023 | 11,205,906 | C:98.68%[S:96.56%,D:2.12%],<br>F:0.61%,I:0.00%,M:0.71% |
| *Agrypnia vestita* | GCA_016648135.1 | Olsen et al. 2021 | 111,757 | C:91.25%[S:83.10%,D:8.15%],<br>F:4.14%,I:0.09%,M:4.52% |
| *Eubasilissa regina* | GCA_022840565.1 | Kawahara et al. 2022 | 29,378,647 | C:98.73%[S:96.28%,D:2.45%],<br>F:0.80%,I:0.00%,M:0.47% |
| *Nectopsyche paramo* | JAWQED000000000 | New | 1,090,281 | C:97.31%[S:95.24%,D:2.07%],<br>F:0.80%,I:0.00%,M:1.88% |
| *Athripsodes cinereus* | GCA_947579605.1 | Darwin Tree of Life Project | 948,465 | C:97.22%[S:96.61%,D:0.61%],<br>F:1.08%,I:0.00%,M:1.69% |
| *Odontocerum albicorne* | GCA_949825065.1 | Darwin Tree of Life Project | 387,033 | C:97.22%[S:96.75%,D:0.47%],<br>F:1.18%,I:0.00%,M:1.60% |
| *Limnocentropus insolitus* | Available on FigShare | New | 33,230,923 | C:98.87%[S:98.26%,D:0.61%],<br>F:0.56%,I:0.00%,M:0.56% |

## Results

### Opsin Distribution and Gene Tree

The number of opsin paralogs found within each species ranged from three to as many as nine (Fig. 3). Opsin diversity within Integripalpia (Fig. 3: Clade 2) was more variable than that of Annulipalpia (Fig. 3: Clade 1). For example, the number of LW opsins among species of Integripalpia ranged from one to five paralogs. While we recovered all three visual opsins in four of the seven species within “basal Integripalpia” (Fig. 3: Clade 2-a), the three remaining species from the family Rhyacophilidae were found to have lost the SW opsin. Conversely, in the tube-case making Integripalpia (Fig. 3: Clade 2-b), the SW opsin was lost in all except *Nectopsyche paramo* and *Athripsodes cinereus*, both of which are from the family Leptoceridae.

The opsin sequences formed four distinct clades in the gene tree corresponding to the LW, SW, UV, and RH7 opsin groups (Fig. 2). Within each opsin clade, the arrangements of suborders were maintained, but occasionally interspecific relationships of opsin sequences did not match those in the established species tree (Frandsen et al., 2024). We only found minor differences between the CDS and peptide gene trees, primarily within the LW and UV opsin groups (Fig. S2). However, none of these areas of incongruence occurred across opsin classes and they were in areas of the trees with lower bootstrap support values (Fig. S2). The LW opsins formed two distinct clades in the opsin gene tree, a phenomenon observed in other insect orders (Feuda et al., 2016; Futahashi et al., 2015; Guignard et al., 2022; Lord et al., 2016; Mulhair et al. 2023; Sharkey et al., 2017; Sharkey et al., 2023; Sondhi et al., 2021, Suvorov et al., 2017). All species had at least one LW opsin in the LW1 clade, while only 14 species—distributed across both suborders—had an additional LW2 opsin (Fig. S1).

**Figure 2:**
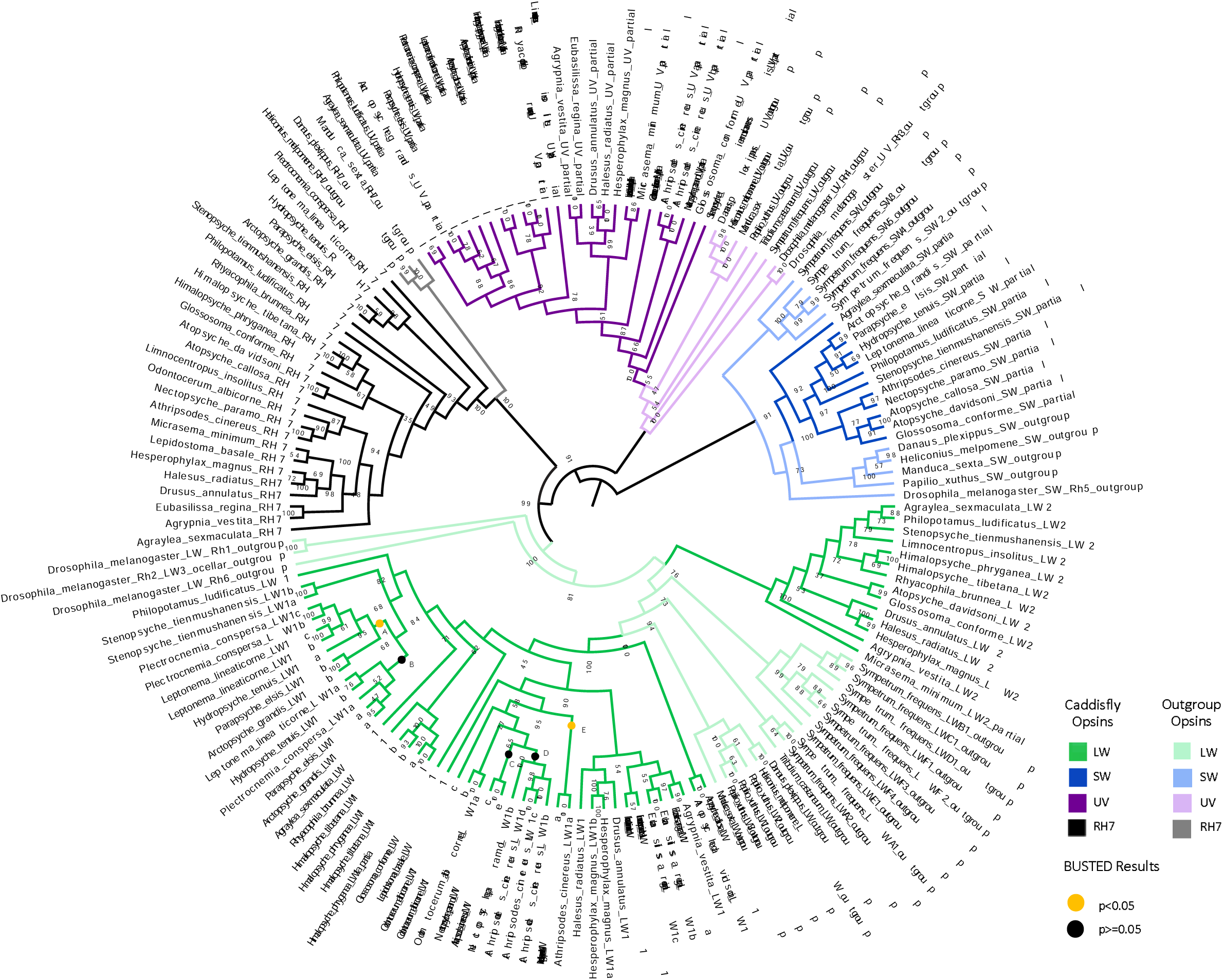
Maximum likelihood tree from caddisfly and outgroup opsin DNA sequences. Nodes are labeled with bootstrap values and branches are colored by opsin type. Letters at the end of the node labels (e.g., LW1<u>a</u>, LW1<u>b</u>, etc.) denote multiple copies of that opsin type in a species. Clades marked with a dot (A-E) were tested for episodic diversifying selection with BUSTED (Table S3).

### Lineage-Specific Gene Duplication Events

We observed a few instances of paralogs that were paired in the opsin gene tree (Fig. 2) and adjacent to each other on the same contig in the genome assembly (Table S1), suggesting independent tandem duplication events.. This was true for copies of the UV opsin in *Athripsodes cinereus* and the LW1 opsin in *Stenopsyche tienmushanensis*, *Leptonema lineaticorne*, *Plectrocnemia conspersa*, *Hesperophylax magnus*, *Eubasilissa regina*, *Odontocerum albicorne*, and *Athripsodes cinereus* (Fig. 2). Upon testing the branches of these duplicate opsins for selection with the Branch-Site Unrestricted Statistical Test for Episodic Diversification (BUSTED, Kosakovsky Pond et al., 2020; Murrell et al., 2015), we found evidence of episodic diversifying selection for three opsin paralogs: *Leptonema lineaticorne* LW1c, *Stenopsyche tienmushanensis* LW1b, and *Eubasilissa regina* LW1b (Table S3).

We also found instances of gene duplication in the common ancestor of closely related species. For example, Clades A and B in Fig. 2 each have one or more LW1 opsin from the species *Leptonema lineaticorne*, *Hydropsyche tenuis*, *Parapsyche elsis*, *Arctopsyche grandis*, and *Plectrocnemia conspersa*, all but the latter of which belong to the family Hydropsychidae. We tested for episodic diversifying selection using BUSTED (Kosakovsky Pond et al., 2020; Murrell et al., 2015) and found evidence of selection in the sequences in Clade A but not Clade B (Fig, 2, Table S3). Similarly, multiple duplication events occurred in the LW1 opsins in the lineage leading to *Nectopsyche paramo* and *Athripsodes cinereus*, both from the family Leptoceridae (Fig. 2: Clades C, D, and E). Among these duplications, we only found evidence of selection in the opsin sequences belonging to Clade E (Table S3).

## Discussion

We searched across 25 caddisfly genome assemblies to determine the number and phylogenetic relationships of opsins in Trichoptera. Our results suggest that caddisfly opsin evolution is likely driven by life history strategies and ambient light conditions as found in other insect orders (Briscoe et al., 2010; Feuda et al., 2016; French et al., 2015, Futahashi et al., 2015; Guignard et al., 2022; Lord et al., 2016; Mulhair et al., 2023; Sharkey et al., 2017; Sondhi et al., 2021; Suvorov et al., 2017).

We found some incongruencies between the gene tree and the species tree. Given the relatively smaller number of characters compared to the reference species tree, which was generated from genome-wide data (Frandsen et al., 2024), this is not unexpected and could be due to stochastic error from under-sampling of characters. This is also evidenced by lower bootstrap values in areas of the tree that were incongruent with the species tree (Fig. 2).

The distribution of opsins within Annulipalpia, the fixed-retreat makers, was relatively invariable. Interestingly, their ecological distributions are also less varied; most species inhabit fast-moving streams as larvae and are short-lived in the riparian zone as adults. While there were a few LW1 duplications in this suborder, the only loss we observed was the SW opsin gene in *Plectrocnemia conspersa* (Fig. 3). When searching for opsin genes, we found a sequence highly similar to the SW opsin in this species; however, we excluded it from the dataset due to the presence of stop codons. Given that the genome assembly of *Plectrocnemia conspersa* was high quality, we hypothesize that this is likely a true loss (Table 1).

**Figure 3:**
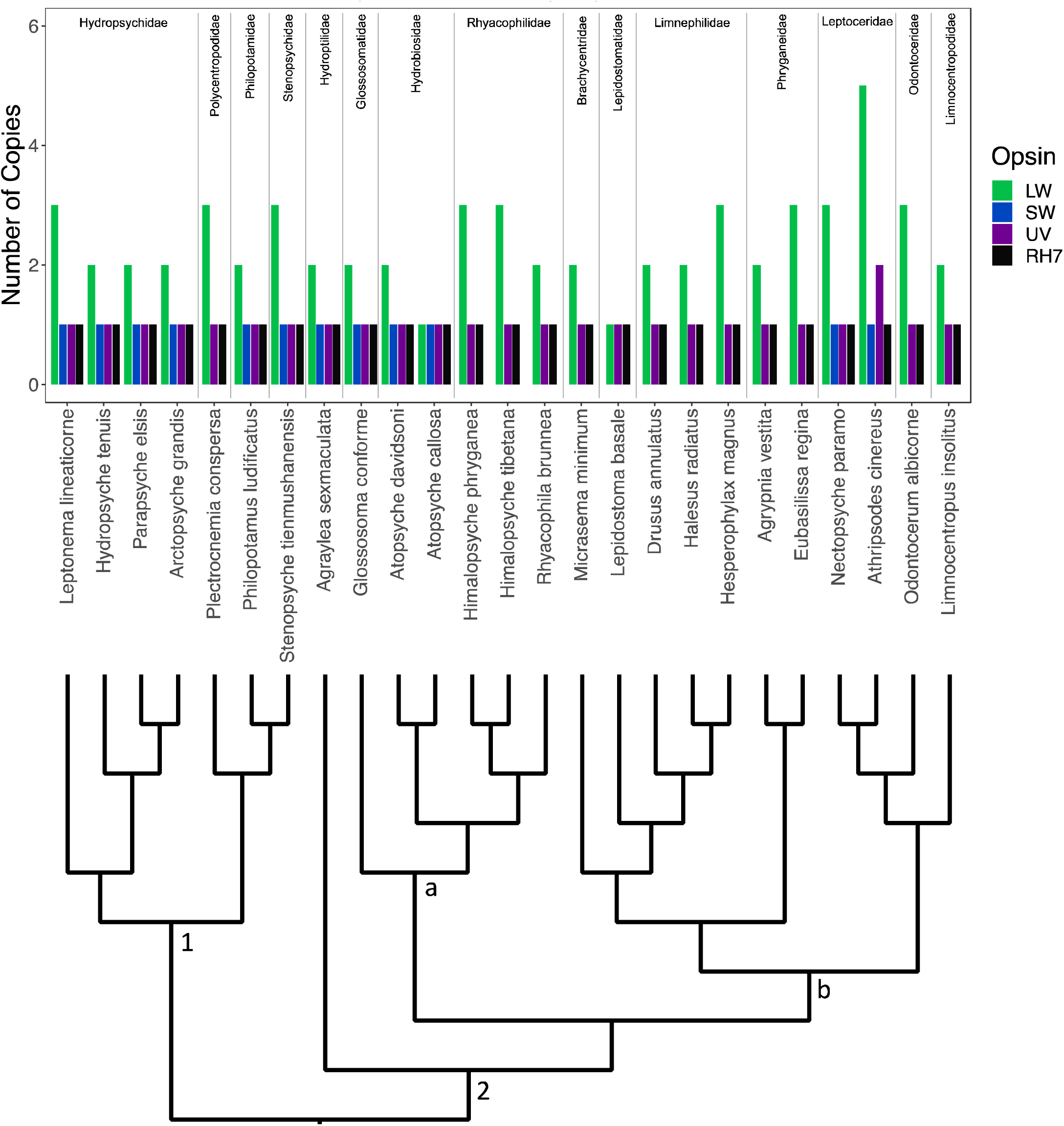
Bar plot of the number of opsin copies in each caddisfly genome. The bars are colored by opsin type. The species are ordered by the species phylogeny indicated below the bar plot, which is based on a recent study that examined the caddisfly phylogeny in depth (Frandsen et al., 2024). Families are labeled above the bar plot. Suborders are labeled within the phylogeny as follows: (1) Annulipalpia—retreat-makers; (2) Integripalpia—cocoon– and tube case-makers; (a) (and *Agraylea sexmaculata*) basal Integripalpia—cocoon-makers; (b) tube case making Integripalpia.

The SW opsin gene was also lost in Integripalpia within the basal cocoon-making family, Rhyacophilidae, and within the majority of the clade of tube-case makers (Fig. 3 Clade 2-b). Loss of the SW opsin in other groups such as the American cockroach (French et al., 2015) and Neuropteroidea (Lord et al., 2016; Sharkey et al., 2017; Sharkey et al., 2023) is hypothesized to be associated with the low-light environments in which the ancestors of these insects lived. Caddisflies are primarily crepuscular and thus are most active during low-light conditions, which may be related to the loss of the SW opsin in Rhyacophilidae and in many tube-case making caddisflies.

Interestingly, in contrast to most species within the tube-case makers, we were able to find the SW opsin in both *Nectopsyche paramo* and *Athripsodes cinereus.* The latter was also the only species with two UV opsins and the highest number of LW opsins. *Nectopsyche paramo* and *Athripsodes cinereus* both belong to the family Leptoceridae, a family with known sexual eye dimorphism (Gullefors & Petersson, 1993), long adult antennae, and many genera with intricate wing patterns and brightly colored or iridescent wing hairs and scales (Fig.1)(Holzenthal et al., 2007). The high number of opsins and presence of the SW opsin could be related to the variety in wing coloration and patterns in some species of Leptoceridae. Future work should combine more sampling within this interesting area of the caddisfly phylogeny with gene expression and physiological data to better model the visual system of Trichoptera and to test hypotheses related to color vision and opsin diversity.

To further investigate the role of the duplication events we observed in the LW and UV opsin groups, we tested paralogs in these areas of the tree for positive selection. We found evidence of episodic diversifying selection in some LW1 opsin paralogs but did not detect evidence of selection in the UV opsin paralogs (Table S3). In each instance when a paralog was found to be under selection, the duplicate paralog was not found to be under selection, possibly suggesting a route to neofunctionalization in those copies undergoing diversifying selection. Recent work in Lepidoptera and Hemiptera has identified instances of family– and species-specific duplications of visual opsins leading to adaptations that extend visual capacity (Briscoe et al., 2010; Feuda et al., 2016; Finkbeiner & Briscoe, 2021; Frentiu et al., 2007; Friedrich 2023; Mulhair et al., 2023; Sison-Mangus et al., 2008; Spaethe & Briscoe, 2004). Denser taxon sampling in future work can help clarify the evolutionary timing of duplication events and the mechanisms and role of selection that we uncovered.

Here, we conducted the first comprehensive study of visual opsins in Trichoptera. We found that the species with the highest diversity of opsins were derived from the group with known sexual eye dimorphism (Gullefors & Petersson, 1993) and, which also contains species of the most colorful and intricately patterned wings, the Leptoceridae. Opsin evolution in caddisflies also may have been driven by life-history strategies and the low light conditions during which caddisflies are active. The findings of this study provide a basis for future research on the diverse and complex visual systems in Trichoptera.

## Materials and Methods

We assessed 25 species of Trichoptera using five newly assembled genomes (details in Supplemental Note S1), and 20 publicly available genome assemblies (Table 1). Using the 1000 opsin sequences found by Guignard et al. (2022), we performed a tBLASTn search of opsin sequences against each caddisfly genome, keeping hits with an e-value < 10^-40^. The resulting opsin sequences were extracted from their corresponding genomes using Pyfaidx (Shirley et al., 2015). We filtered redundant hits from multiple queries by extracting the widest window from each contig and then classifying the gene phylogenetically downstream.

We performed gene prediction using AUGUSTUS v3.4.0 (Stanke, et al. 2006) followed by a BLASTp search against the online NCBI databases, maintaining only hits similar to other insect opsins. We then manually checked the annotations in Geneious Prime v2023.0.4 (https://www.geneious.com), using outgroup sequences as a guide, to ensure that the entire gene was correctly annotated (see Supplementary Note S1 for more details). We included opsin sequences from a variety of insect orders for outgroup comparison: Lepidoptera (*Danaus plexippus, Heliconius melpomene, Manduca sexta, Papilio xuthus*), Odonata (*Sympetrum frequens)*, Diptera (*Drosophila melanogaster*), and Coleoptera (*Tribolium castaneum*), all accessed through GenBank (Benson et al., 2013). We performed additional searches for opsin genes to further verify the absence of the SW and LW2 opsins in many species (see Supplementary Note S1 for more details). We also provide supplementary tables (Table S2a,b,c) with information on the completeness of each visual opsin sequence.

### Opsin Gene Tree Reconstruction

To determine phylogenetic relationships among opsin sequences, we first aligned the opsin peptide sequences using MAFFT v7.487 (Katoh & Standley, 2013) and created a codon alignment with PAL2NAL v14.1 (Suyama, 2009). We performed phylogenetic reconstruction on both the CDS and peptide alignments by first selecting the best substitution model using ModelFinder (Kalyaanamoorthy et al., 2017; Minh et al., 2020), then performing a maximum likelihood tree search with 1000 UltraFast bootstrap replicates corrected with the bootstrap nearest neighbor interchange option enabled to guard against the risk of overestimating bootstrap support (-bb 1000 –bnni). We viewed the resulting trees in FigTree v1.4.4 (Rambaut, 2018). To highlight differences between the CDS and peptide trees, we created a face-to-face comparison in R with ggtree v3.2.1 (Yu et al., 2016) and ggplot2 v3.3.5 (Wickham, 2016) (Fig. S2).

### Selection Analysis

To assess the duplications in the LW and UV opsin groups, we created separate codon alignments for both the LW and UV opsin groups with MAFFT v7.487 (Katoh & Standley, 2013) and PAL2NAL v14.1 (Suyama, 2009), then tested for episodic diversifying selection using BUSTED as implemented in HyPhy (Kosakovsky Pond et al., 2020; Murrell et al., 2015). We tested branches of species-specific duplicated opsins individually as well as five deeper duplication events (Fig. 2: Clades A-E) and reported the resulting p-values in Table S3.

### Adult Caddisfly Illustrations

### Opsin Gene Tree

### Opsin Counts by Species

### Genome Assemblies

## Acknowledgements

We thank the BYU Office of Research Computing for the compute resources that we used in this study. We thank the BYU College of Life Sciences for providing funding through the College Undergraduate Research Award Program. We thank Ed Wilcox from the BYU DNA Sequencing Center for sequencing the new genomes presented here. J.H. acknowledges funding from the Deutsche Forschungsgemeinschaft, Project number 502865717, B.R.-T., S.U.P., R.W.H., and P.B.F. acknowledge funding from the Dirección General de Investigación, Universidad de Las Américas (Ecuador): “Montane freshwater diversity, from taxonomy to functional genomics, an approximation from Trichoptera-Part II.” (AMB.BRT.23.02).

## Authors contributions

Conceptualization: A.P. and P.B.F. Formal analysis: A.P., J.H., E.R.G., and P.B.F. Resources: J.H., B.R.-T., R.B.K., and P.B.F. Data curation: A.P., J.H., and E.R.G. Writing-original draft prep: A.P. Writing-review and editing: all authors. Visualization: A.P., and R.W.H. Supervision: P.B.F. Funding acquisition: A.P., J.H., S.U.P., B.R.-T., and P.B.F.

