## Supplemental Information for "Evolution of Opsin Genes in Caddisflies (Insecta: Trichoptera)"

**Supplementary Information**

**Supplementary Note S1: Additional Materials and Methods**

*Genome Assembly*

We provide new genomes for the following five species: *Atopsyche callosa* (Navás, 1924), *Limnocentropus insolitus* Ulmer, 1907, *Nectopsyche paramo* Holzenthal & Ríos-Touma, 2018, *Plectrocnemia conspersa* (Curtis, 1834), and *Rhyacophila brunnea* Banks, 1911. We extracted DNA from the head and thorax of a single individual for each species using a Qiagen Genomic-tip extraction kit. Next, we used pulse-field gel electrophoresis to visualize high molecular weight DNA. The DNA for each extraction was then sheared to 15 kbp fragments using the Diagenode Megaruptor and PacBio libraries were generated using the SMRTbell Express Template Prep Kit 2.0. Each library was run on 30-hour 8M PacBio SMRT cells on the Sequel II instrument at the Brigham Young University DNA Sequencing Center. Then, using PacBio SMRTlink software, we computed HiFi reads (with quality above Q20) from the raw data. Lastly, we assembled the raw reads into assemblies using HiCanu and Hifiasm v.0.13-r307 (Cheng et al., 2021, Nurk et al. 2020). Genome quality was assessed using compleasm v.0.2.2 using the Endopterygota OrthoDB v.10 gene set (Huang & Li, 2023) (Table 1).

The *Nectopsyche paramo* assembly was of lower quality than the others due to low sequence coverage. We further improved this assembly by first additional haplotig purging with purge_dups (Guan et al., 2020). For this, we first ran minimap2 (Li et al, 2018) with options -I6G and -xmap-pb to align the reads to the genome and generate a paf file which was used to calculate read depth histogram and base-level read depth with pbcstat as follows: pbcstat *fastq.paf.gz && calcuts PB.stat > cutoffs 2>calcults.log. We then split the assembly using split_fa and did a self-self alignment with minimap2 using parameters -I6G -xasm5 -DP. Haplotigs and overlaps were purged with purge_dups using the following command: purge_dups -2 -T cutoffs -c PB.base.cov $pri_asm.split.self.paf.gz > dups.bed 2 > purge_dups.log. Finally, the purged primary and haplotig sequences from the draft assembly were obtained with get_seqs using the dups.bed file. We also performed an additional scaffolding step by "rescuing" subreads which did not contribute to building HiFi reads. For this, we kept the longest subread per polymerase read that was not collapsed into a HiFi using custom perl scripts. We used these reads to scaffold the contigs from the purged draft assembly using Sspace longreads 1-1 (Boetzer, 2014). To close gaps in the scaffolded assembly, we use tgs gaploser 1.0.1 (Xu et al., 2020) with the HiFi reads using parameters --tgstype pb and --ne. Since the HiFi reads have been used for the draft assembly, they usually do not fall into the gaps. For this reason, we conducted another round of gapclosing with tgs gapcloser 1.0.1 and racon/1.4.3 (Vaser et al., 2017) with the longest subreads per polymerase read.

We screened the final genome assembly for potential contamination with taxon-annotated GC-coverage (TAGC) plots using BlobTools v1.1.1 (Laetsch et al., 2017). For this purpose, the bam file resulting from the back-mapping analysis was converted to a BlobTools readable.cov file with “blobtools map2cov”. Taxonomic assignment for BlobTools was done with BLASTn 2.10.0+ (Camacho et al., 2009) using -task megablast and -e-value 1e-25. The blobDB was created and plotted from the cov file and blast hits. Contamination detected by NCBI were filtered as follows: scaffold1292: prok:b-proteobacteria; scaffold291:253360..253403:adaptor:NGB00972.1; scaffold413:405475..405519, 405779..405823, 407187..407270: adaptor:NGB00972.1.

*Manual Gene Annotation*

To manually verify the annotation for each opsin sequence, we first aligned the translation—from the AUGUSTUS annotation—with the outgroup sequences of the same opsin type using MUSCLE (Edgar, 2004). In instances where the caddisfly opsin sequence did not fully align with the outgroup sequences, the annotation was manually modified in Geneious Prime v2023.0.4 (<https://www.geneious.com>) until the entire gene sequence aligned with the outgroups. This was done by first removing or shortening CDS regions that did not align with or varied greatly from the outgroup sequences, causing large gaps in the alignment. Then, for regions of the gene that were present in the outgroup sequences but missing from the annotation, we used the search tool in Geneious to find conserved motifs from within those regions and added or extended CDS annotations accordingly. When extending and shortening the CDS regions of the annotation, we ensured that every intron region occurred at canonical splice sites. After we made these manual adjustments to the annotation, we re-aligned the translation with the outgroup sequences, and the alignment was reassessed for completeness, i.e. that all exons and core motif regions were present in the alignment for each sequence from each species. If additional changes were necessary, the same process of adjusting the annotation was followed until the gene aligned with the outgroup sequences.

*SW and LW2 Gene Loss*

After annotating all the opsins genes found from the BLAST search, we performed an additional tBLASTn search against the genomes using the orthologous SW and LW2 sequences that were successfully obtained from the other caddisfly species. We maintained BLAST hits that were in regions of the genomes we had not previously checked, and we followed the same steps for annotating the gene as described previously. Additionally, we further checked for the SW opsin in some species using synteny of BUSCO genes. To do this we found the BUSCO genes flanking the SW gene from a caddisfly genome that contained the SW opsin. We then located the area with the same BUSCO genes in the genomes with the putative gene loss and searched for the SW opsin both with BLAST and manually in Geneious.

*Gene Completeness*

We report the completeness of each visual opsin gene in supplementary Tables S2a, b, and c). We used the number and identity of exons in Lepidoptera as a guide in making the tables, i.e. the column labels “Exon 1, Exon 2, …” refer to the exons present in Lepidoptera. Within the SW and UV opsin groups, the first exon in the lepidopteran outgroup sequences was not highly conserved, making it difficult to recover this exon in the caddisfly opsins despite searching for it both in Geneious and with BLAST. Therefore, these sequences are labeled as partial sequences in the opsin gene trees. In addition, there were a few instances where two exons were combined into a single exon in the caddisfly sequences. For example, the fifth and sixth exons in the lepidopteran UV opsin formed a single continuous exon, with no intron, in the caddisfly UV opsin.

*Long Wavelength Opsin Counts by Species*

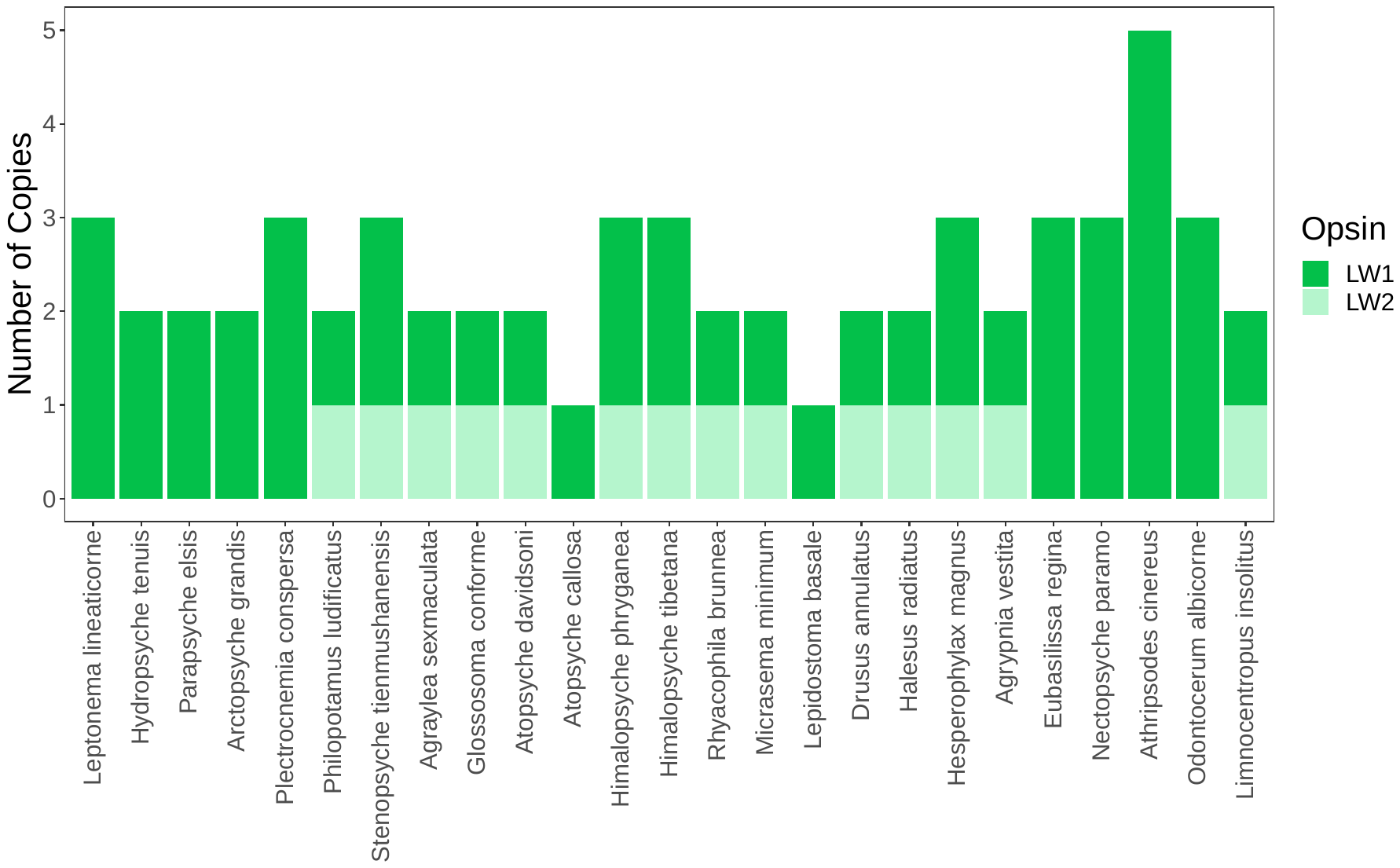

*Figure S1:* Bar plot of the number of LW opsin sequences in each caddisfly genome. Different shades of green are used to distinguish between which LW clade (LW1 vs LW2) the sequence belongs to in the opsin gene tree.

*CDS vs Peptide Opsin Gene Tree*

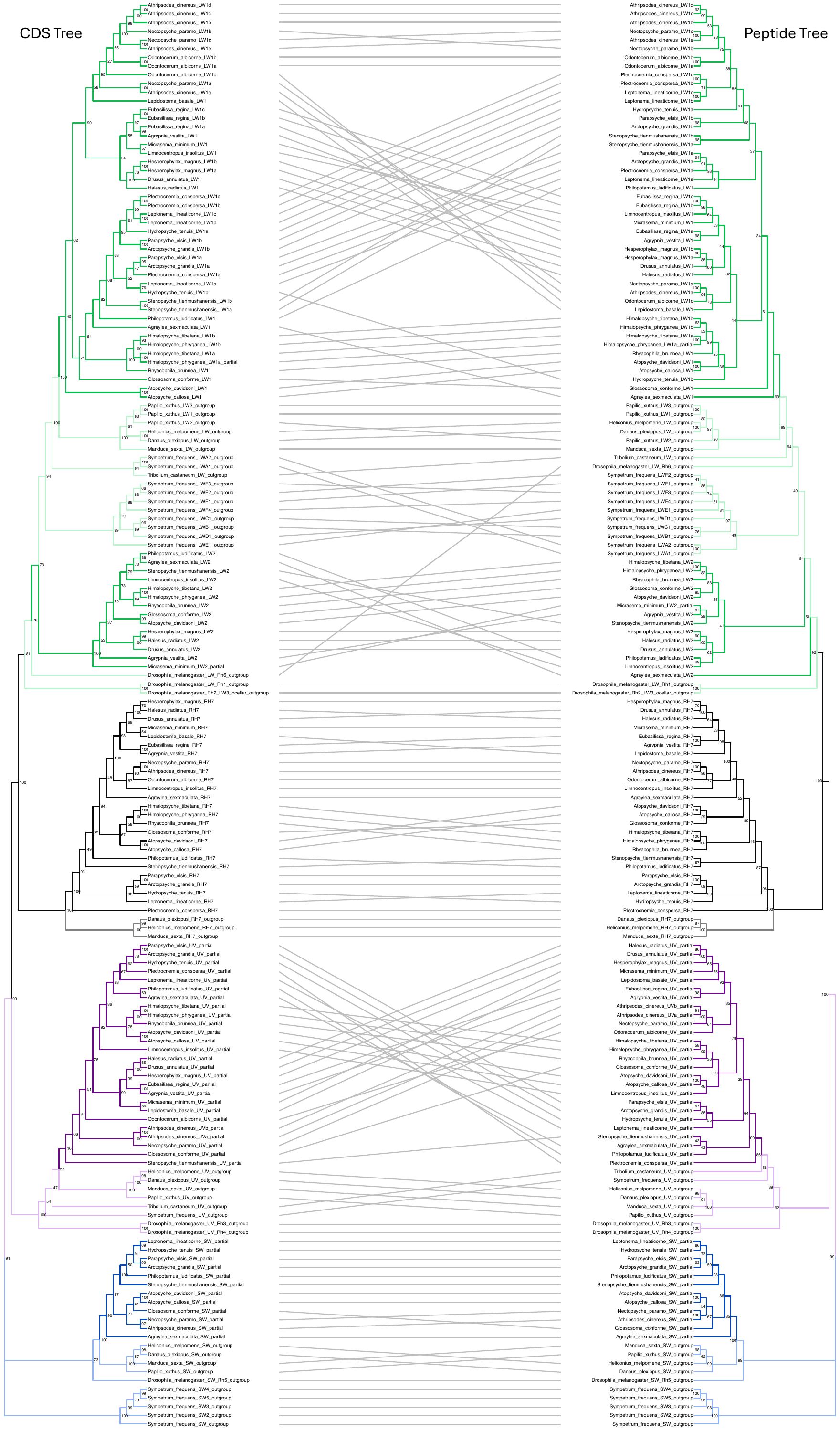

*Figure S2*: Face-to-face comparison of opsin gene trees. The tree on the left was constructed from the CDS alignment, and the tree on the right was constructed from the amino acid alignment. Lines between the trees connect branches for the same opsin sequence. Nodes are labeled with bootstrap values and branches are colored by opsin type. Letters at the end of the node labels (e.g., LW1a, LW1b, etc.) denote multiple copies of that opsin type in a species.

*Opsin Copies by Contig/Scaffold*

| **Species** | **Contig** | **Opsins** | **Count** |
| --- | --- | --- | --- |
| Leptonema lineaticorne | ptg000022l | LW1a,LW1b,LW1c | 3 |
| Leptonema lineaticorne | ptg000007l | UV | 1 |
| Leptonema lineaticorne | ptg000016l | RH7 | 1 |
| Leptonema lineaticorne | ptg000010l | SW | 1 |
| Hydropsyche tenuis | VTON01000025.1 | LW1a,LW1b | 2 |
| Hydropsyche tenuis | VTON01000015.1 | UV | 1 |
| Hydropsyche tenuis | VTON01000103.1 | RH7 | 1 |
| Hydropsyche tenuis | VTON01000016.1 | SW | 1 |
| Parapsyche elsis | ctg4 | LW1a,LW1b | 2 |
| Parapsyche elsis | ctg10 | UV | 1 |
| Parapsyche elsis | ctg2 | RH7 | 1 |
| Parapsyche elsis | ctg26 | SW | 1 |
| Arctopsyche grandis | ptg000036l | LW1a,LW1b | 2 |
| Arctopsyche grandis | ptg000003l | UV | 1 |
| Arctopsyche grandis | ptg000066l | RH7 | 1 |
| Arctopsyche grandis | ptg000027l | SW | 1 |
| Plectrocnemia conspersa | ptg000010l | LW1a,LW1b,LW1c | 3 |
| Plectrocnemia conspersa | ptg000014l | UV,RH7 | 2 |
| Philopotamus ludificatus | 36677 | LW1,LW2 | 2 |
| Philopotamus ludificatus | 31889 | UV | 1 |
| Philopotamus ludificatus | 37236 | RH7 | 1 |
| Philopotamus ludificatus | 35719 | SW | 1 |
| Stenopsyche tienmushanensis | WACJ01000159.1 | LW1a,LW1b | 2 |
| Stenopsyche tienmushanensis | WACJ01000038.1 | LW2 | 1 |
| Stenopsyche tienmushanensis | WACJ01000469.1 | UV | 1 |
| Stenopsyche tienmushanensis | WACJ01000088.1 | RH7 | 1 |
| Stenopsyche tienmushanensis | WACJ01000174.1 | SW | 1 |
| Agraylea sexmaculata | ctg2218 | LW1,LW2 | 2 |
| Agraylea sexmaculata | ctg4359 | UV | 1 |
| Agraylea sexmaculata | ctg3042 | RH7 | 1 |
| Agraylea sexmaculata | ctg5594 | SW | 1 |
| Glossosoma conforme | ctg4 | LW1,LW2 | 2 |
| Glossosoma conforme | ctg233 | UV | 1 |
| Glossosoma conforme | ctg100 | RH7 | 1 |
| Glossosoma conforme | ctg77 | SW | 1 |
| Atopsyche davidsoni | ptg000024l | LW1 | 1 |
| Atopsyche davidsoni | ptg000018l | LW2 | 1 |
| Atopsyche davidsoni | ptg000023l | UV,SW | 2 |
| Atopsyche davidsoni | ptg000021l | RH7 | 1 |
| Atopsyche callosa | ptg000005l | LW1 | 1 |
| Atopsyche callosa | ptg000011l | UV,SW | 2 |
| Atopsyche callosa | ptg000026l | RH7 | 1 |
| Himalopsyche phryganea | ctg43 | LW1a,LW1b,LW2 | 3 |
| Himalopsyche phryganea | ctg64 | UV | 1 |
| Himalopsyche phryganea | ctg36 | RH7 | 1 |
| Himalopsyche tibetana | ptg000052l | LW1a,LW1b,LW2 | 3 |
| Himalopsyche tibetana | ptg000013l | UV | 1 |
| Himalopsyche tibetana | ptg000023l | RH7 | 1 |
| Rhyacophila brunnea | ptg000092l | LW1,LW2 | 2 |
| Rhyacophila brunnea | ptg001293l | UV | 1 |
| Rhyacophila brunnea | ptg000497l | RH7 | 1 |
| Micrasema minimum | ctg549 | LW1,LW2 | 2 |
| Micrasema minimum | ctg851 | UV | 1 |
| Micrasema minimum | ctg97 | RH7 | 1 |
| Lepidostoma basale | scf586 | LW1 | 1 |
| Lepidostoma basale | scf1634 | UV | 1 |
| Lepidostoma basale | scf536 | RH7 | 1 |
| Drusus anulatus | ctg183 | LW1,LW2 | 2 |
| Drusus anulatus | ctg639 | UV | 1 |
| Drusus anulatus | ctg533 | RH7 | 1 |
| Halesus radiatus | scf12671 | LW1,LW2 | 2 |
| Halesus radiatus | scf12849 | UV | 1 |
| Halesus radiatus | scf9559 | RH7 | 1 |
| Hesperophylax magnus | JAIUSX010000031.1 | LW1a,LW1b,LW2 | 3 |
| Hesperophylax magnus | JAIUSX010000029.1 | UV | 1 |
| Hesperophylax magnus | JAIUSX010000014.1 | RH7 | 1 |
| Agrypnia vestita | JADDOH010006403.1 | LW1,LW2a | 2 |
| Agrypnia vestita | JADDOH010024855.1 | LW2b | 1 |
| Agrypnia vestita | JADDOH010023784.1 | UV | 1 |
| Agrypnia vestita | JADDOH010000163.1 | RH7 | 1 |
| Eubasilissa regina | tig00000571 | LW1a,LW1b,LW1c | 3 |
| Eubasilissa regina | tig00000026 | UV | 1 |
| Eubasilissa regina | tig00000001 | RH7 | 1 |
| Nectopsyche paramo | scaffold202 | LW1a,LW1b,LW1c | 3 |
| Nectopsyche paramo | scaffold70 | UV | 1 |
| Nectopsyche paramo | scaffold4 | RH7 | 1 |
| Nectopsyche paramo | scaffold102 | SW | 1 |
| Athripsodes cinereus | OX388326.1 | LW1a,LW1b,LW1c,LW1d,LW1e | 5 |
| Athripsodes cinereus | OX388314.1 | UVa,UVb,SW | 3 |
| Athripsodes cinereus | OX388315.1 | RH7 | 1 |
| Odontocerum albicorne | OX463860.1 | LW1a,LW1b,LW1c | 3 |
| Odontocerum albicorne | OX463845.1 | UV | 1 |
| Odontocerum albicorne | OX463846.1 | RH7 | 1 |
| Limnocentropus insolitus | ptg000030l | LW1,LW2 | 2 |
| Limnocentropus insolitus | ptg000008l | UV | 1 |
| Limnocentropus insolitus | ptg000018l | RH7 | 1 |

*Table S1:* Table showing which opsin genes are on each contig/scaffold of the genome assembly for each species.

*Completeness of Long Wavelength Opsins by Exon*

| **Opsin** | **Exon 1** | **Exon 2** | **Exon 3** | **Exon 4** | **Exon 5** | **Exon 6** | **Exon 7** | **Exon 8** |
| --- | --- | --- | --- | --- | --- | --- | --- | --- |
| Agraylea sexmaculata LW1 | C | C | C | C | C | C | C | C |
| Agraylea sexmaculata LW2 | C | C | C | C | C | C | C | C |
| Agrypnia vestita LW1 | C | C | C | C | C | C | C | C |
| Agrypnia vestita LW2 | C | C | C | C | C | C | C | C |
| Arctopsyche grandis LW1a | C | C | C | C | C | C | C | C |
| Arctopsyche grandis LW1b | C | C | C | C | C | C | C | C |
| Athripsodes cinereus LW1a | C | C | C | C | C | C | C | C |
| Athripsodes cinereus LW1b | C | C | C | C | C | C | C | C |
| Athripsodes cinereus LW1c | C | C | C | C | C | C | C | C |
| Athripsodes cinereus LW1d | C | C | C | C | C | C | C | C |
| Athripsodes cinereus LW1e | C | C | C | C | C | C | C | C |
| Atopsyche callosa LW1 | C | C | C | C | C | C | C | C |
| Atopsyche davidsoni LW1 | C | C | C | C | C | C | C | C |
| Atopsyche davidsoni LW2 | C | C | C | C | C | C | C | C |
| Drusus annulatus LW1 | C | C | C | C | C | C | C | C |
| Drusus annulatus LW2 | C | C | C | C | C | C | C | C |
| Eubasilissa regina LW1a | C | C | C | C | C | C | C | C |
| Eubasilissa regina LW1b | C | C | C | C | C | C | C | C |
| Eubasilissa regina LW1c | C | C | C | C | C | C | C | C |
| Glossosoma conforme LW1 | C | C | C | C | C | C | C | C |
| Glossosoma conforme LW2 | C | C | C | C | C | C | C | C |
| Halesus radiatus LW1 | C | C | C | C | C | C | C | C |
| Halesus radiatus LW2 | C | C | C | C | C | C | C | C |
| Hesperophylax magnus LW1a | C | C | C | C | C | C | C | C |
| Hesperophylax magnus LW1b | C | C | C | C | C | C | C | C |
| Hesperophylax magnus LW2 | C | C | C | C | C | C | C | C |
| Himalopsyche phryganea LW1a | P | C | C | C | C | C | C | C |
| Himalopsyche phryganea LW1b | C | C | C | C | C | C | C | C |
| Himalopsyche phryganea LW2 | C | C | C | C | C | C | C | C |
| Himalopsyche tibetana LW1a | C | C | C | C | C | C | C | C |
| Himalopsyche tibetana LW1b | C | C | C | C | C | C | C | C |
| Himalopsyche tibetana LW2 | C | C | C | C | C | C | C | C |
| Hydropsyche tenuis LW1a | C | C | C | C | C | C | C | C |
| Hydropsyche tenuis LW1b | C | C | C | C | C | C | C | C |
| Lepidostoma basale LW1 | C | C | C | C | C | C | C | C |
| Leptonema lineaticorne LW1a | C | C | C | C | C | C | C | C |
| Leptonema lineaticorne LW1b | C | C | C | C | C | C | C | C |
| Leptonema lineaticorne LW1c | C | C | C | C | C | C | C | C |
| Limnocentropus insolitus LW1 | C | C | C | C | C | C | C | C |
| Limnocentropus insolitus LW2 | C | C | C | C | C | C | C | C |
| Micrasema minimum LW1 | C | C | C | C | C | C | C | C |
| Micrasema minimum LW2 | C | P | C | C | C | C | C | C |
| Nectopsyche paramo LW1a | C | C | C | C | C | C | C | C |
| Nectopsyche paramo LW1b | C | C | C | C | C | C | C | C |
| Nectopsyche paramo LW1c | C | C | C | C | C | C | C | C |
| Odontocerum albicorne LW1a | C | C | C | C | C | C | C | C |
| Odontocerum albicorne LW1b | C | C | C | C | C | C | C | C |
| Odontocerum albicorne LW1c | C | C | C | C | C | C | C | C |
| Parapsyche elsis LW1a | C | C | C | C | C | C | C | C |
| Parapsyche elsis LW1b | C | C | C | C | C | C | C | C |
| Philopotamus ludificatus LW1 | C | C | C | C | C | C | C | C |
| Philopotamus ludificatus LW2 | C | C | C | C | C | C | C | C |
| Plectrocnemia conspersa LW1a | C | C | C | C | C | C | C | C |
| Plectrocnemia conspersa LW1b | C | C | C | C | C | C | C | C |
| Plectrocnemia conspersa LW1c | C | C | C | C | C | C | C | C |
| Rhyacophila brunnea LW1 | C | C | C | C | C | C | C | C |
| Rhyacophila brunnea LW2 | C | C | C | C | C | C | C | C |
| Stenopsyche tienmushanensis LW1a | C | C | C | C | C | C | C | C |
| Stenopsyche tienmushanensis LW1b | C | C | C | C | C | C | C | C |
| Stenopsyche tienmushanensis LW2 | C | C | C | C | C | C | C | C |

*Table S2a*: Table showing the completeness of each long wavelength opsin in this study. Columns represent exons in lepidopteran outgroups. Labels indicate if each exon is complete (C), partial (P), or missing (M).

*Completeness of Short Wavelength Opsins by Exon*

| **Opsin** | **Exon 1** | **Exon 2** | **Exon 3** | **Exon 4** | **Exon 5** | **Exon 6** | **Exon 7** | **Exon 8** |
| --- | --- | --- | --- | --- | --- | --- | --- | --- |
| Agraylea sexmaculata SW | M | P | C | C | C | C | C | C |
| Arctopsyche grandis SW | M | P | C | C | C | C | C | C |
| Athripsodes cinereus SW | M | P | C | C | C | C | C | C |
| Atopsyche callosa SW | M | P | C | C | C | C | C | C |
| Atopsyche davidsoni SW | M | P | C | C | C | C | C | C |
| Glossosoma conforme SW | M | P | C | C | C | C | C | C |
| Hydropsyche tenuis SW | M | P | C | C | C | C | C | C |
| Leptonema lineaticorne SW | M | P | C | C | C | C | C | C |
| Nectopsyche paramo SW | M | P | C | C | C | C | C | C |
| Parapsyche elsis SW | M | P | C | C | C | C | C | C |
| Philopotamus ludificatus SW | M | P | C | C | C | C | C | C |

*Table S2b*: Table showing the completeness of each short wavelength opsin in this study. Columns represent exons in lepidopteran outgroups. Labels indicate if each exon is complete (C), partial (P), or missing (M).

*Completeness of Ultraviolet Wavelength Opsins by Exon*

| **Opsin** | **Exon 1** | **Exon 2** | **Exon 3** | **Exon 4** | **Exon 5** | **Exon 6** | **Exon 7** | **Exon 8** |
| --- | --- | --- | --- | --- | --- | --- | --- | --- |
| Agraylea sexmaculata UV | M | P | C | C | C | C | C | C |
| Agrypnia vestita UV | M | P | C | C | C | C | C | C |
| Arctopsyche grandis UV | M | P | C | C | C | C | C | C |
| Athripsodes cinereus UVa | M | P | C | C | C | C | C | C |
| Athripsodes cinereus UVb | M | P | C | C | C | C | C | C |
| Atopsyche callosa UV | M | P | C | C | C | C | C | C |
| Atopsyche davidsoni UV | M | P | C | C | C | C | C | C |
| Drusus annulatus UV | M | P | C | C | C | C | C | C |
| Eubasilissa regina UV | M | P | C | C | C | C | C | C |
| Glossosoma conforme UV | M | P | C | C | C | C | C | C |
| Halesus radiatus UV | M | P | C | C | C | C | C | C |
| Hesperophylax magnus UV | M | P | C | C | C | C | C | C |
| Himalopsyche phryganea UV | M | P | C | C | C | C | C | C |
| Himalopsyche tibetana UV | M | P | C | C | C | C | C | C |
| Hydropsyche tenuis UV | M | P | C | C | C | C | C | C |
| Lepidostoma basale UV | M | P | C | C | C | C | C | C |
| Leptonema lineaticorne UV | M | P | C | C | C | C | C | C |
| Limnocentropus insolitus UV | M | P | C | C | C | C | C | C |
| Micrasema minimum UV | M | P | C | C | C | C | C | C |
| Nectopsyche paramo UV | M | P | C | C | C | C | C | C |
| Odontocerum albicorne UV | M | P | C | C | C | C | C | C |
| Parapsyche elsis UV | M | P | C | C | C | C | C | C |
| Philopotamus ludificatus UV | M | M | M | C | C | C | C | C |
| Plectrocnemia conspersa UV | M | P | C | C | C | C | C | C |
| Rhyacophila brunnea UV | M | P | C | C | C | C | C | C |
| Stenopsyche tienmushanensis UV | M | P | C | C | C | C | C | C |

*Table S2c*: Table showing the completeness of each UV opsin in this study. Columns represent exons in lepidopteran outgroups. Labels indicate if each exon is complete (C), partial (P), or missing (M).

*Selection Analysis*

| **Branch/Clade** | **BUSTED P-value** |
| --- | --- |
| Leptonema lineaticorne LW1b | 0.5 |
| Leptonema lineaticorne LW1c | 0.04955 |
| Plectrocnemia conspersa LW1b | 0.2014 |
| Plectrocnemia conspersa LW1c | 0.5 |
| Stenopsyche tienmushanensis LW1a | 0.5 |
| Stenopsyche tienmushanensis LW1b | 0.04291 |
| Athripsodes cinereus LW1b | 0.5 |
| Athripsodes cinereus LW1c | 0.5 |
| Odontocerum albicorne LW1a | 0.5 |
| Odontocerum albicorne LW1b | 0.5 |
| Odontocerum albicorne LW1c | 0.5 |
| Eubasilissa regina LW1b | 0.04513 |
| Eubasilissa regina LW1c | 0.5 |
| Hesperophylax magnus LW1b | 0.5 |
| Athripsodes cinereus UVa | 0.4993 |
| Athripsodes cinereus UVb | 0.5 |
| Clade A | 0.0008536 |
| Clade B | 0.3858 |
| Clade C | 0.3581 |
| Clade D | 0.1003 |
| Clade E | 0.01382 |

*Table S3*: Results from running BUSTED on various branches and clades in the opsin CDS gene tree. Clades that were tested are labeled in the opsin gene tree (Fig. 2).
